# Extraction and Characterization of Rice Bran Polysaccharides and Their Derivative Preparation and Antioxidant Activity

**DOI:** 10.1101/2020.11.06.370965

**Authors:** Gangliang Huang

## Abstract

The crude polysaccharide of rice bran polysaccharide was extracted with water from defatted rice bran, and the optimal reaction conditions of the experiment were determined. The reaction temperature is 100°C, the reaction time is 5h, the solid-liquid ratio is 1:10; the concentration ratio of polysaccharide to Sevag reagent is 2:1, and rice polysaccharide is precipitated by ethanol insoluble property and ethanol. The ethanol concentration ratio is 3:1. Using the methods and conditions of this experiment, the maximum polysaccharide yield is 2.18%. Then the chemical structure of rice bran polysaccharide was analyzed by IR, UV and 13CNMR methods. Three rice bran polysaccharide derivatives were prepared and characterized: carboxymethylated rice bran polysaccharide, phosphorylated rice bran polysaccharide and acetylated rice bran polysaccharide. Also studied the antioxidant activity of rice bran polysaccharide and its derivatives (hydroxy radical ion scavenging ability, superoxide anion scavenging ability, anti-lipid peroxidation ability, reducing ability), and found that phosphorylated rice bran polysaccharide has good activity. The acetylated rice bran polysaccharide may weaken the original activity.

## 1 Introduction

Rice is a member of the grass family Gramineae and is the second largest food crop in the world. Global rice production accounts for about 37% of total grain output. At present, the annual global rice output has reached about 500 million tons, and it is still increasing year by year. China’s rice planting area accounts for 1/4 of the country’s total grain planting area. The average annual total rice output is more than 200 million tons. Rice bran is a by-product of rice processing, which accounts for about 6% to 7.5% of rice quality.[1] In China, 1 300-1 400 t of rice bran can be produced every year.

Rice bran polysaccharide (Rice bran polysaccharide (RBPs)) is a heteroglycan composed of α-1,3-glucan as the main chain and a variety of monosaccharide units such as xylose, galactose, mannose, rhamnose, arabinose, and ribose as side chains. [2–5]

Rice bran is rich in a variety of active ingredients, such as fats, proteins, carbohydrates, oryzanol, vitamins, calcium, iron, magnesium, manganese, potassium, sodium, zinc and other essential nutrients for the human body.[6–9] Rice bran can be used to extract rice bran oil, make rice wax, rice bran, etc. These products have the effect of lowering blood cholesterol. Rice bran has replaced corn and other raw materials in animal and poultry feed, and there are many studies and reports on reducing feed cost and improving economic benefits. Rice bran has the functions of intestinal tract, appetizer, and qi, and can be used to treat beriberi, cancer, etc. [11–12].

A variety of biological polysaccharides have biological activity with development potential. Their functions include immunomodulation, anti-tumor, hypoglycemic, hypolipidemic, and anti-radiation. Plant polysaccharide research has attracted more and more attention. The international scientific community even proposes that the 21st century is a polysaccharide century Scientific experiment research, antibacterial and antiviral, liver protection and other health effects. Therefore, plant polysaccharides have long been widely used in the field of public life such as the medical community, the catering community. [13–16]

Rice bran polysaccharide also has significant biological activity and health care functions. Studies at home and abroad have shown that rice bran polysaccharides have various active functions such as anti-tumor, hypoglycemic, cholesterol-lowering, anti-bacterial infection and immunity enhancement [17–19]. Unfortunately, at present, most rice bran has not been effectively used, because the lipase activity in rice bran is very high, which is easy to cause rice bran spoilage. Therefore, improving the extraction process of rice bran polysaccharides is of great significance to the utilization of rice bran.

## 2 Materials

For rice bran, we chose the 2019 fresh autumn defatted rice bran provided by Hubei Fuhua Company.This batch of rice bran will be used within four months.

## 3 Methods and Results

### 3.1 Methods

Basic process of extraction and separation and purification of defatted rice bran polysaccharides:

#### 3.1.1 Extraction, Separation and Purification of Rice Bran Polysaccharide

100g of defatted rice bran has been weighed for the extraction of rice bran polysaccharides. The defatted rice bran and distilled water have been placed in a three-necked flask at a feed-water ratio of 1:10, and a magnetic stir bar has been placed in the mixture. After the defatted rice bran has been soaked in distilled water for 1 hour, the whole system was placed in a constant temperature water bath at 100 ° C. The first extraction has been carried out after stirring and heating at 100 ° C for 5 hours, centrifugation at 4000 r/min for 10 min, and the supernatant has been collected. In the second extraction, 200 mL of distilled water has been added to the filter residue, it has been stirred and heated at 100 ° C. for 1 hour, it has been centrifuged again at 4000 r/min for 10 min, and the supernatant is collected. In the third extraction, 200 mL of distilled water has been added to the remaining filter residue, which has been stirred and heated at 100 ° C. for 1 hour, and has been centrifuged at 4000 r/min for 10 min to collect the supernatant. All the collected supernatants that have been extracted three times are combined and concentrated to 200 mL of solution. The rice bran polysaccharide solution was sealed and stored in a refrigerator at 0-5 ° C.

The Sevag reagent method was used to remove protein from rice bran polysaccharides. According to the literature [20], the concentrated crude rice bran polysaccharide solution and Sevag reagent were mixed in a ratio of 2:1. At this time, the protein removal rate is the highest.

200mL of rice bran crude polysaccharide solution, 100mL of Sevag reagent was used to remove protein. The Sevag reagent in our experiment used CHCl_3_ : CH_3_(CH_2_)_3_OH was 2: 1. It should be noted here that we have configured 120mL of Sevag reagent,80mL of CHCl_3_ and 40mL of CH_3_(CH_2_)_3_OH. 100mL Sevag reagent was mixed with 200ml the crude rice bran polysaccharide concentrated solution. After the mixing was completed, a magnetic stirrer was used to stir for 30 minutes, in order to fully react the crude polysaccharide solution with Sevag reagent. After the reaction, it was allowed to stand for 30 minutes to separate the layers, and then centrifuged at 4000 r/min for 10 min. The solution has been divided into three layers. The supernatant has been transferred into the beaker with a dropper, and the middle and lower solution are poured into the waste bottle for recovery. Then the supernatant was repeated the above operation twice to remove protein.

Taking advantage of the insoluble properties of rice bran polysaccharides, 95% ethanol was used to precipitate rice bran polysaccharides. After reading the reference[21–22], we know that 3 times the volume of 95% ethanol is used to precipitate rice bran polysaccharides. The ethanol was stirred for 5 minutes to make it contact completely, and after standing for 5 hours, a light yellow flocculent precipitate was obtained, which was centrifuged at 4000 r/min for 10 min. The bottom sediment was rice bran polysaccharide. The mortar containing the rice bran polysaccharide was placed in a vacuum freeze dryer for 12 h, and the dried rice bran polysaccharide solid was ground into a powder. The weight of the rice bran polysaccharide powder was weighed and placed in a plastic tube. Waiting to be used.

#### 3.1.2 Determination of Polysaccharide Content

The polysaccharide content was determined by the phenol sulfuric acid method [23].Preparation of standard curve: According to [24], a 100 mg/L glucose standard solution was prepared. Different volumes of glucose standard solutions were drawn into test tubes and filled with deionized water to 0.5ml(0.5ml deionized water was used as the blank control group).0.50 mL of 5% phenol solution was added and mixed thoroughly, 2.5 mL of concentrated sulfuric acid was added and quickly shaken, and cooled at room temperature for 15 minutes to room temperature.Distilled water was used as a blank control group. The absorbance of glucose solutions with different mass concentrations at 490 nm (A) has been measured. The mass concentration of glucose has been taken as the abscissa and the absorbance has been taken as the ordinate. A standard curve has been drawn, and the regression equation has been calculated. The regression equation of **Fig.2** is: Y = 13.082x-0.0091,R2 =0.997. Finally, three parallel experiments are performed on each sample. The polysaccharide content can be calculated by the regression equation.

**Fig.1.**
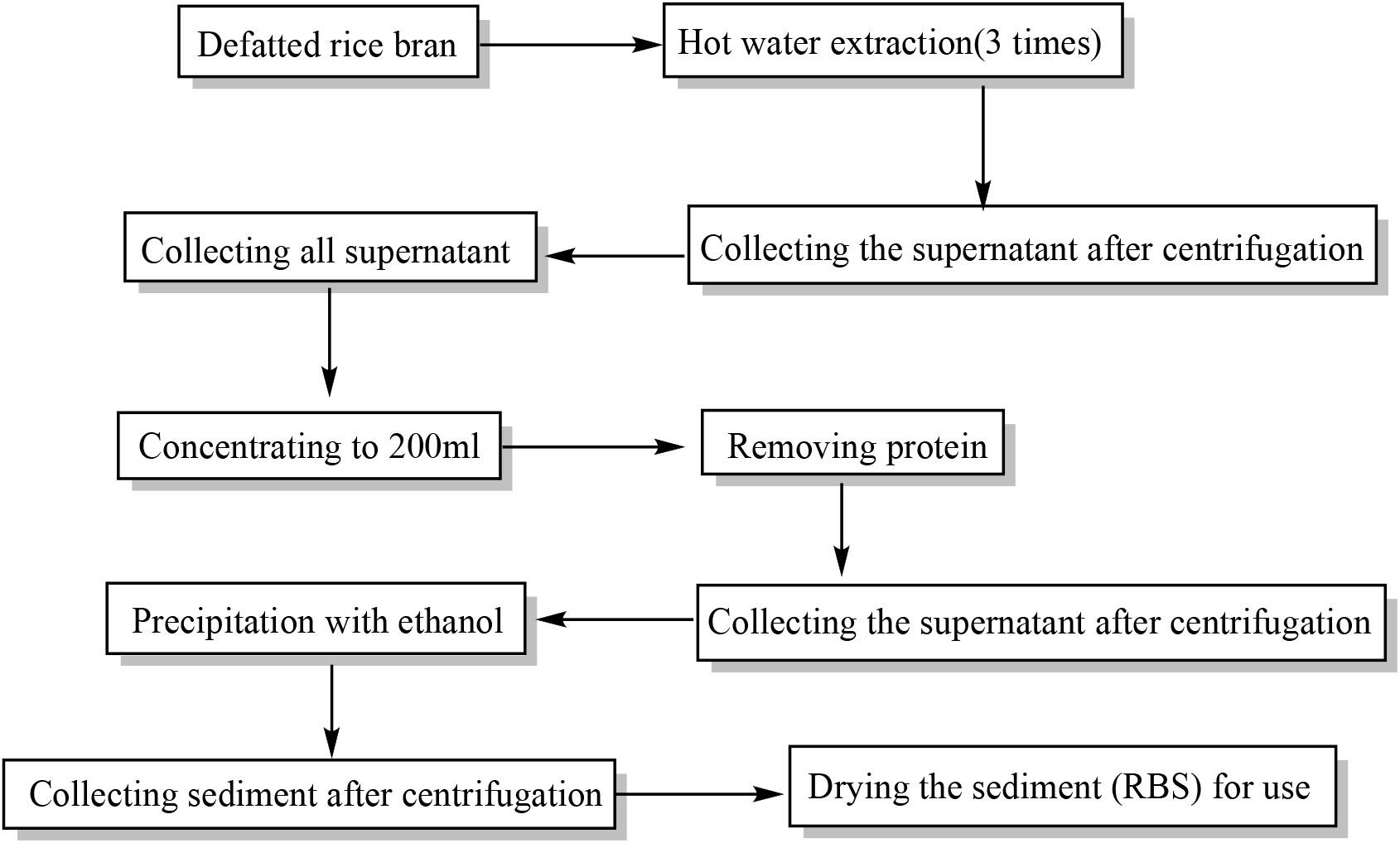
Rice bran polysaccharide preparation process

**Fig.2.**
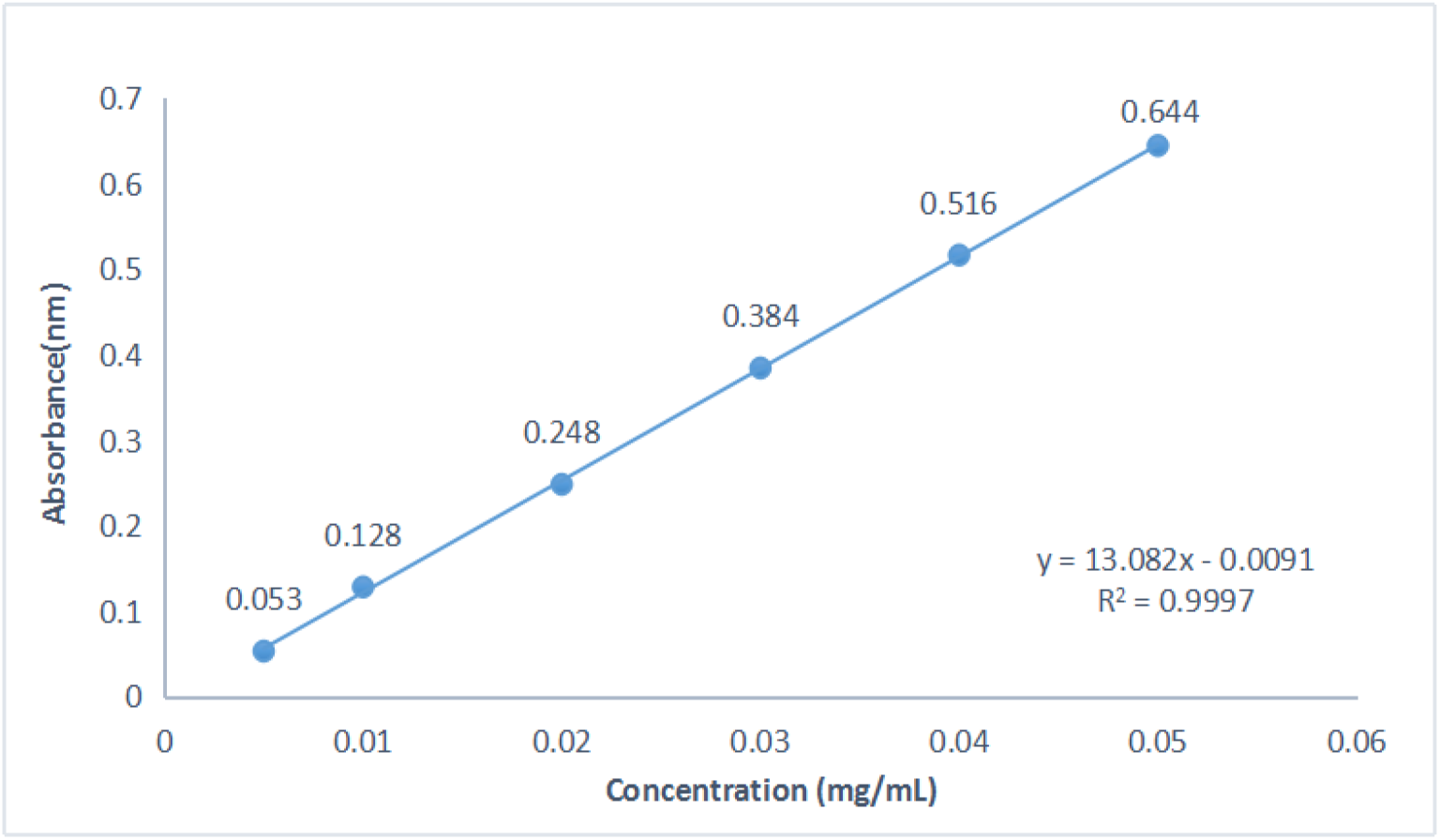
The standard curve

#### 3.1.3 Preparation of carboxymethylated rice bran polysaccharide [25]

Carboxymethylated rice bran polysaccharide.At room temperature, 0.5 g of rice bran polysaccharides (RBPs) was added to 60 ml of isopropanol and stirred for 30 minutes.Then, 30% NaOH solution (5.6 mL) and TCA (2.4 g) were added to the mixture. And the mixture was stirred at 50°C for 3 hours. The reaction solution was adjusted to neutral with dilute hydrochloric acid, and then the dialysis bag was used in the distilled water for 3 days.The supernatantliquid was concentratedto 20 mL. Add 3 times the volume of ethanol, centrifuge, and wash twice with alcohol. The precipitate was dissolved in distilled water and freeze-dried to obtain a white substance (CM-RBPs).

#### 3.1.4 Preparation of phosphorylated rice bran polysaccharide

Phosphorylated rice bran polysaccharide.RBPs (0.5 g) was suspended in 30 mL DMSO at room temperature with stirring for 8 h. Then adding 6.5 mL pyridine to the mixture, stirring for 0.5 h, and the reaction reagent POCl_3_ (2.5 mL) were added dropwise.After the reaction, the mixture was cooled to room temperature and the pH value was adjusted to 7 with 1 mol/L NaOH solution.The mixtures were precipitated with ethanol (95%), washed and then dialyzed with distilled water for 3d to remove pyridine and salt.The supernatant was concentrated to 20 mL, and add 3 times the volume of ethanol, then centrifuged, washed with alcohol for two times.And then the mixture was dried in a vacuum freezing dryer.Phosphorylated rice bran polysaccharide were collected as P-RBPs.The samples were stored in a desiccator at room temperature till for use.

#### 3.1.5 Preparation of acetylated rice bran polysaccharide

0.5 g of rice bran polysaccharide was dissolved in 10 mL of distilled water, and the pH was adjusted to 9 with NaOH solution (0.5 mol/L). At 30°C, 0.6 ml of acetic anhydride (added multiple times) was added to the polysaccharide solution.After the addition of acetic anhydride has been completed, the reaction was carried out at a constant temperature for 2 hours.After the reaction, the pH of the reaction solution was adjusted to 7 with 1 mol/mL HCl. The reaction solution was filled into a dialysis bag with a molecular weight of 3000 kDa.Distilled water was dialyzed for 48 hours. After the dialysate was concentrated, it was precipitated with 3 times the volume of ethanol (95%) for 24 hours.The precipitate was freeze-dried to obtain acetylated rice bran polysaccharide (Ac-RBPs).

#### 3.1.6 Purification of polysaccharide

The (0.3 g) crude polysaccharide was dissolved in 80 mL distilled water (70 °C, 20 min). The crude polysaccharide was dissolved completely and centrifuged (4000 rpm) with a magnetic stirrer for 8 min. The supernatant was separated by column chromatography on Sephadex G-100, and washed with distilled water. Sulphuric acid-phenol method was used to track and detect the change of sugar content during chromatography.

#### 3.1.7 Discussion on the conditions of water extraction of crude rice bran polysaccharide

Consulting references, it is found that the main factors affecting the extraction rate (E%) of rice bran polysaccharide extracted by water extraction are: reaction temperature (T), reaction time (Time) and solid-liquid ratio (Rs-l).

In order to determine the optimal conditions for extracting rice bran polysaccharides by water extraction, an orthogonal test has been completed and the results are as follows:

① When the reaction temperature is 80° C.
② When the reaction temperature is 100° C.
③ When the reaction temperature is 120° C.

It can be found from the **Table.3** that when the temperature is 80° C, the reaction time is 6h, the solid-liquid ratio 1:15 polysaccharide extraction rate is up to 1.96%.It can be found from the **Table.4** that when the temperature is 100° C, the reaction time is 6h, the solid-liquid ratio 1:15 polysaccharide extraction rate is up to 2.19%.It can be found from the **Table.5** that when the temperature is 120 ° C, the reaction time is 4h, the solid-liquid ratio 1:15 polysaccharide extraction rate ias up to 2.13%.

**Table.3.** Relationship between polysaccharide extraction rate and time and solid-liquid ratio (80° C)

| E% \ Rs-l<br>Time (h) | 1:5 | 1:10 | 1:15 |
| --- | --- | --- | --- |
| 4 | 1.32 | 1.53 | 1.64 |
| 5 | 1.47 | 1.61 | 1.87 |
| 6 | 1.51 | 1.88 | 1.96 |

**Table.4.** Relationship between polysaccharide extraction rate and time and solid-liquid ratio (100° C)

| E% \ Rs-l<br>Time (h) | 1:5 | 1:10 | 1:15 |
| --- | --- | --- | --- |
| 4 | 1.69 | 1.82 | 1.90 |
| 5 | 1.85 | 2.18 | 2.09 |
| 6 | 1.88 | 2.12 | 2.19 |

It can also be found from **Table.4** that when the temperature was at 100 ° C and the reaction time was 5h, the solid-liquid ratio 1:10 polysaccharide extraction rate is 2.18%. While the reaction time was 6h, the solid-liquid ratio 1:15 polysaccharide extraction rate was 2.19%, which was only slightly higher than the former. Therefore, from the perspective of saving energy, the water leaching conditions selected in this experiment were a temperature of 100 ° C, a reaction time of 5h, and a solid-liquid ratio of 1:10.

**Table.4.** Relationship between polysaccharide extraction rate and time and solid-liquid ratio (120° C)

| E%<br>Rs-l<br>Time (h) | 1:5 | 1:10 | 1:15 |
| --- | --- | --- | --- |
| 4 | 1.87 | 2.05 | 2.13 |
| 5 | 1.85 | 1.98 | 2.03 |
| 6 | 1.70 | 1.83 | 1.90 |

#### 3.1.8 Discussion on the conditions for precipitation of rice bran polysaccharides with ethanol

Because the concentration of ethanol and the volume ratio of ethanol to polysaccharide concentrate will affect the yield of rice bran polysaccharide, an orthogonal test was done. Concentration of ethanol (80%, 95%, 100%) volume ratio of ethanol to concentrate (2: 1, 3: 1, 4: 1) Because the concentrate is 200ml, the volume of ethanol is (400ml, 600ml, 800ml).

In **Table.5**, (m) is the weight of the precipitated rice bran crude polysaccharide, (n) is the ethanol concentration, (V) is the volume of ethanol.

**Table.5.** Relationship between the weight of crude polysaccharides from rice bran obtained and the ethanol concentration and ethanol volume

| $m(g)$ \ $V(ml)$ \ $n$ | 400 | 600 | 800 |
| --- | --- | --- | --- |
| 80% | 2.23 | 2.55 | 2.59 |
| 95% | 2.65 | 3.18 | 3.19 |
| 100% | 2.39 | 2.48 | 2.50 |

It can be found from **Table.5** that when the ethanol concentration is 95% and the ethanol volume is 800mL, the yield of crude rice bran polysaccharide is 3.18 g at the maximum. When the ethanol concentration is 95% and the ethanol volume is 600mL, the yield of crude rice bran polysaccharide is slightly lower than the highest value. From the perspective of energy saving, 95% ethanol is used in this experiment, and the volume ratio of ethanol to polysaccharide concentrate is 3: 1.

### 3.2 Results

#### IR-detection

Infrared spectrum detection of rice bran polysaccharide and its derivatives: mixing 3mg with 100mg of dried KBr, grinding, tabletting, and scanning 4000-400 cm-1 wave band by Fourier transform infrared spectrometer.

#### UV-detection

Samples of rice bran polysaccharide and its derivatives were prepared into a 2mg/mL solution, and the 200~600 nm band was scanned by an ultraviolet-visible spectrophotometer (UV-2550).

#### ^13^CNMR

D_2_O was used to prepare a sample solution with a concentration of 40 m g/m l, and the sample solution was analyzed and determined at room temperature by using a 300MHZ full digital nuclear magnetic resonance spectrometer (AVANCE II).

#### 3.2.1 IR detection

It can be found from **Fig.3** that rice bran polysaccharide has a typical infrared absorption peak of polysaccharide. O-H tensile vibration at 3283 cm-1, C-H tensile vibration at 2930 cm^−1^ and C-O-H tensile vibration at 1061 cm^−1^.-CH_2_ symmetric deformation vibration at 1415 cm^−1^, -CH_2_ antisymmetric deformation vibration at 1153.08 cm^−1^, and C-O stretching vibration at 1650.12 cm^−1^.The characteristic absorption peak of saccharide molecular vibration is in the region of 700-950 cm^−1^, and the measurement result has a strong absorption peak at 761 cm^−1^, which is presumed to be caused by the asymmetric contraction of C-O-C skeleton vibration of glucopyranose sugar ring. The absorption peak of 848 cm^−1^ is the characteristic of α-epimer. Therefore, it can be inferred that α-D-glucopyranose is the main component of polysaccharide.

**Fig.3.**
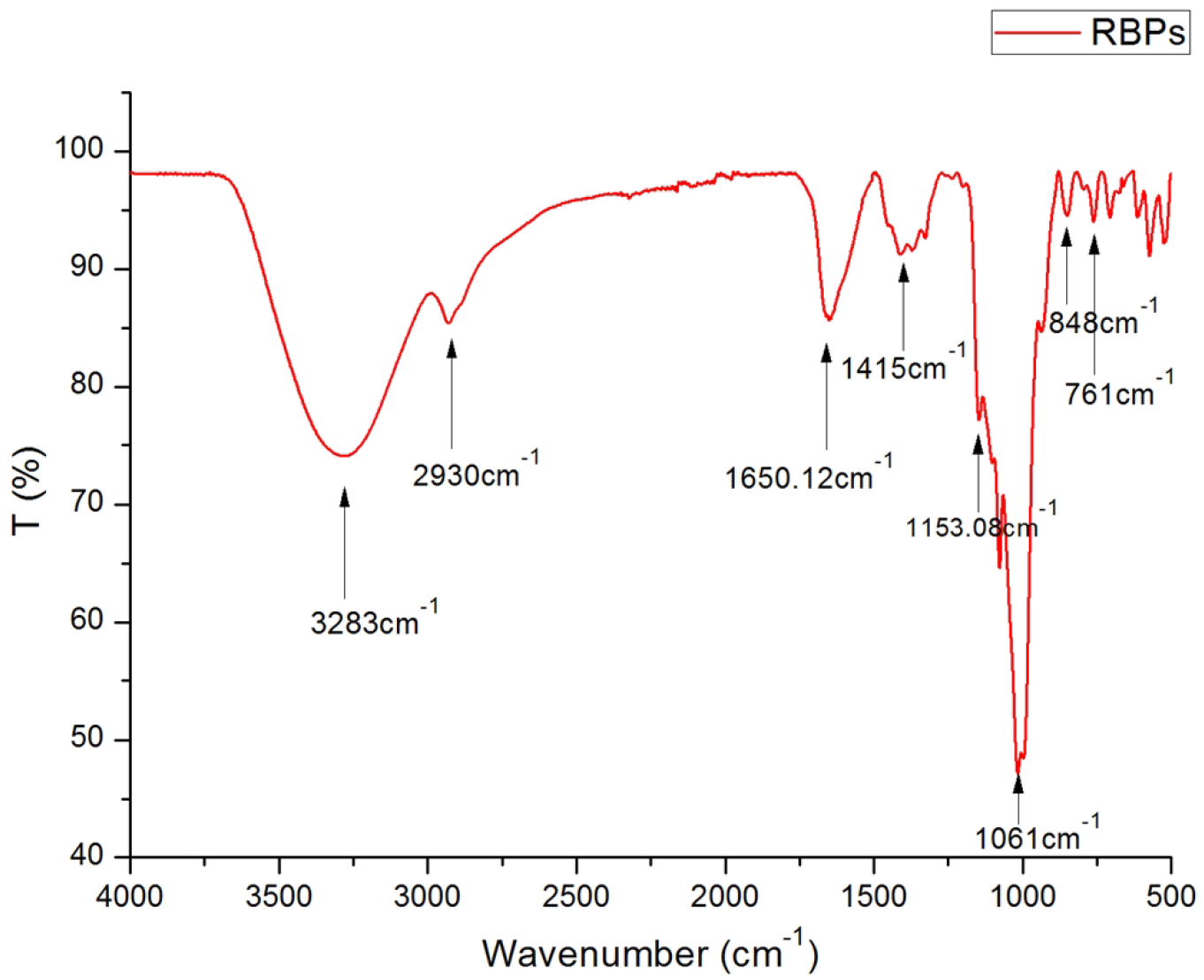
Infrared spectrum of rice bran polysaccharide

From **Fig.4**, we can see that the infrared spectrum of rice bran polysaccharide has no 1603 cm^−1^ and 1329 cm^−1^ (COO-stretch vibration), 1428 cm^−1^ (COO- The bending vibration of the C—H group), these special infrared absorption peaks, meanwhile, they also have absorption peaks typical of rice bran polysaccharides, which can indicate that the experiment of carboxymethylation of rice bran polysaccharides was successful.

**Fig.4.**
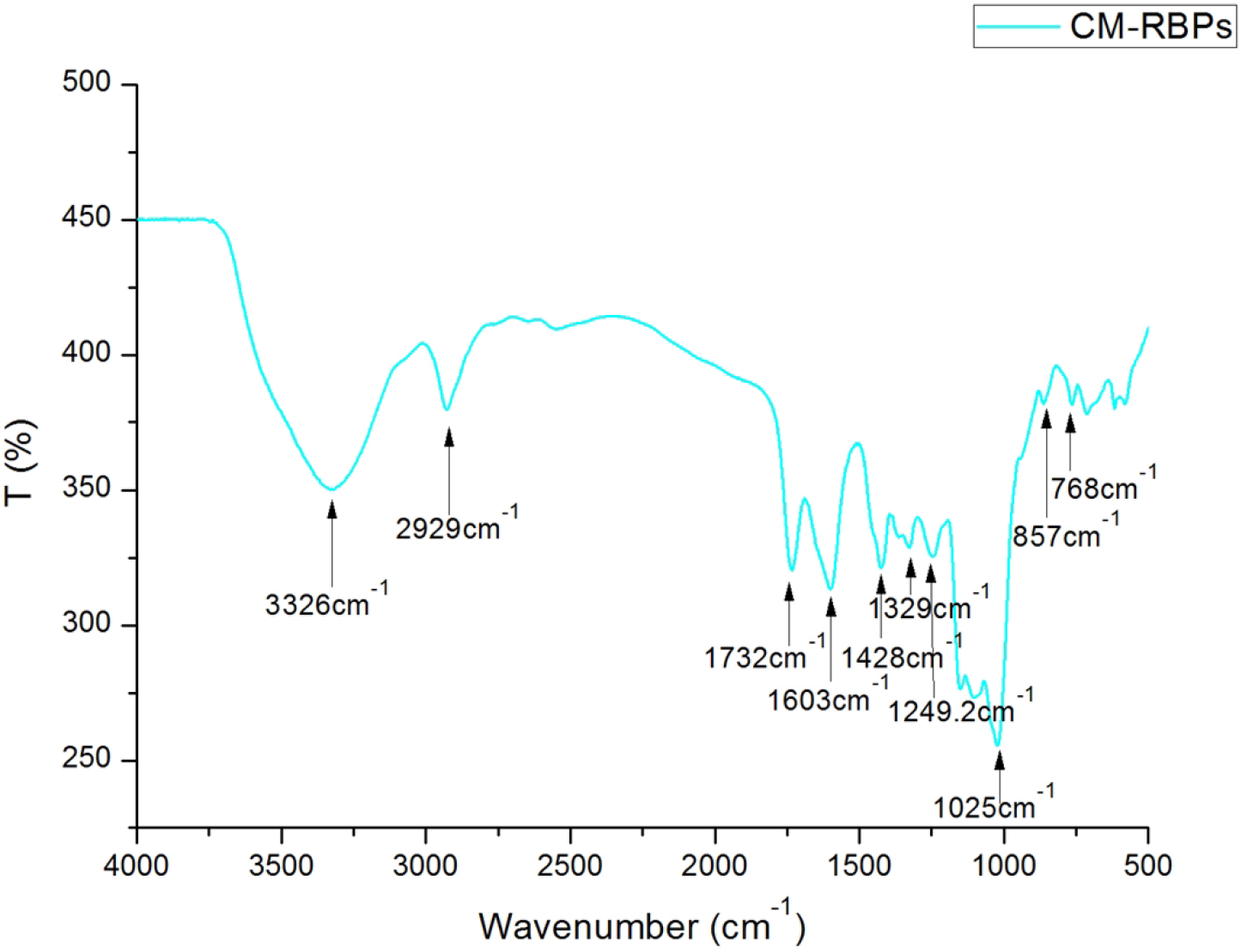
The Infrared spectrum of carboxymethylated rice bran polysaccharide

As shown in **Fig.5**, the infrared spectrum of phosphorylated rice bran polysaccharide ranges from 3000 cm^−1^ to 3600 cm^−1^.At 2933cm^−1^ is the C-H stretching vibration absorption peak in methylene (-CH2-).The broad absorption peak is the characteristic peak of hydroxyl group on sugar ring, the asymmetric stretching peak of C-O at 1632 cm^−1^, the angular vibration of C-H at 1491 cm^−1^, the angular vibration of O-H at 1102 cm^−1^, and a new weak absorption peak at 1264cm^−1^, which is caused by P=O in phosphate. Because the degree of substitution of phosphorylation is not high, and his absorption peak is relatively weak, we suspect that it is caused by hydrolysis of phosphorus oxychloride, which can preliminarily indicate that phosphate is likely to be successfully introduced.

**Fig. 5.**
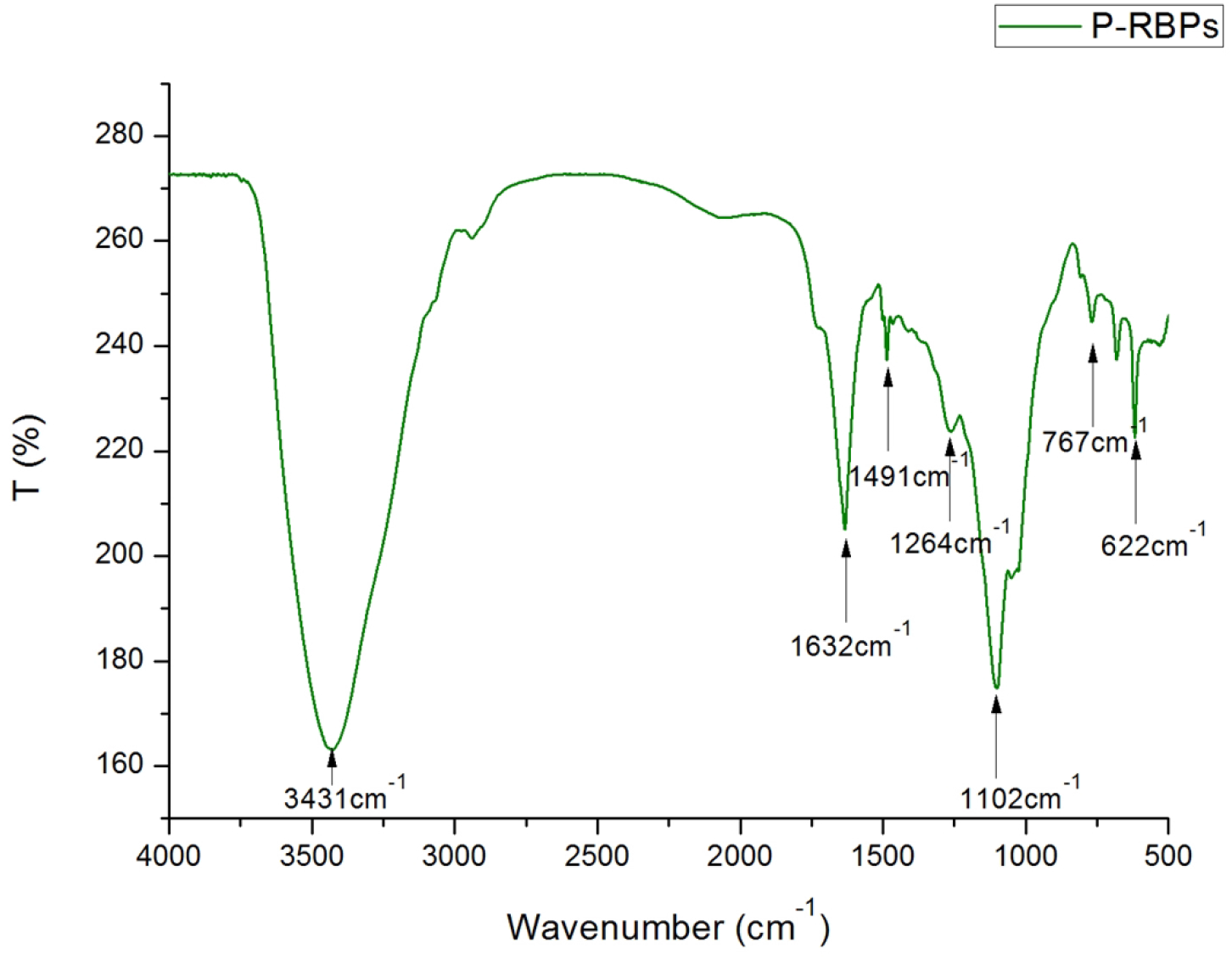
Infrared spectrum of phosphorylated rice bran polysaccharide

**Fig. 6.**
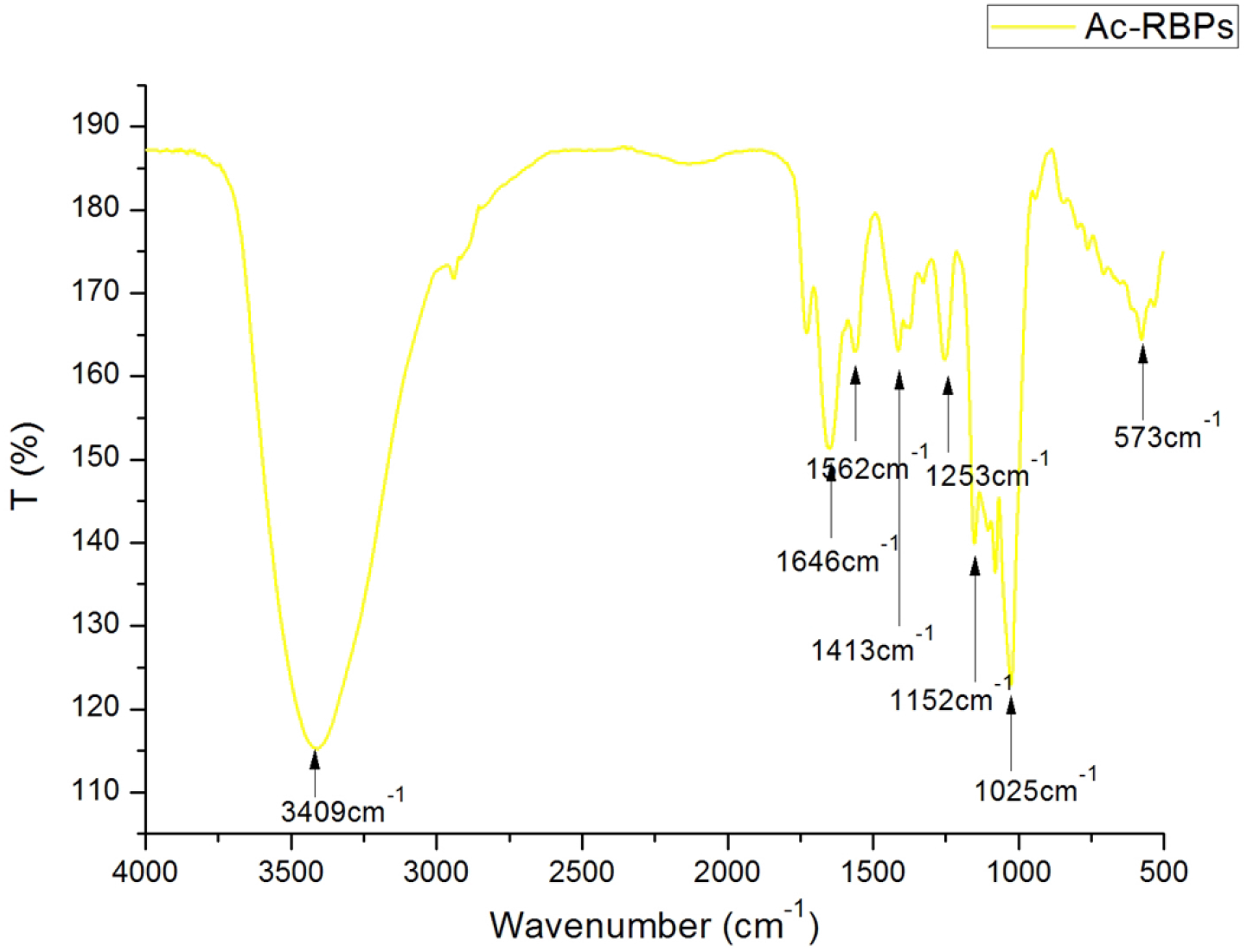
Infrared spectrum of acetylated rice bran polysaccharide

**Fig. 7.**
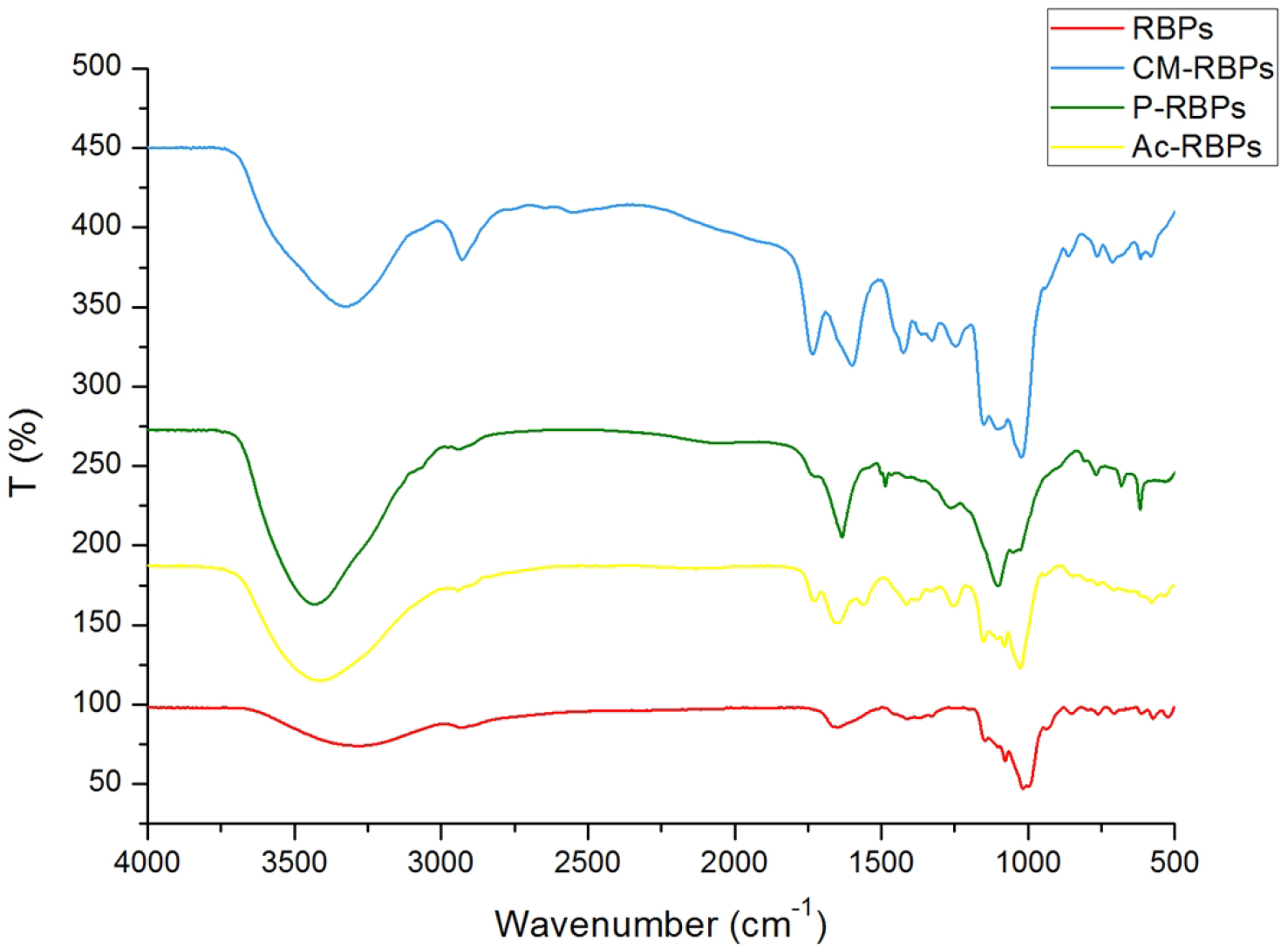
Comparison of infrared spectra of rice bran polysaccharides and several derivatives

At 3409cm^−1^ is the stretching vibration peak of the intramolecular or intermolecular hydrogen bond O-H of the polysaccharide.Ac-RBPs obtained after acetylation modification showed a new absorption peak at 1727cm^−1^ which was the contraction vibration peak of the ester group C=O double bond, and at the same time, a weaker ester group C-O contraction vibration peak appeared at 1253cm^−1^. It shows that the acetyl group has been successfully introduced into the rice bran polysaccha.

#### 3.2.2 UV detection

Rice bran polysaccharide showed a weak absorption peak at 260-280 nm, and the protein had a characteristic absorption peak around 260 nm. The benzene ring of the tyrosine and tryptophan residues in the protein contained conjugated double bonds, so the protein solution was. There is an absorption ultraviolet absorption peak at 280 nm. When the sample contains nucleic acids such as purines and pyrimidines that absorb ultraviolet light, there will be greater interference when measuring the protein content at 280 nm. It indicates that the rice bran polysaccharide may contain a small amount of tyrosine, tryptophan and nucleic acid.[29]

The rice bran polysaccharide has been deproteinized. According to **Fig. 8**, it can be seen that the absorbance of rice bran polysaccharide without deproteinization treatment is highest at 260nm and 280nm, while the absorbance at 260nm and 280nm decreases with the increase of the number of deproteinization. After the three deproteinization treatments, the absorbance decreased significantly.

**Fig.8.**
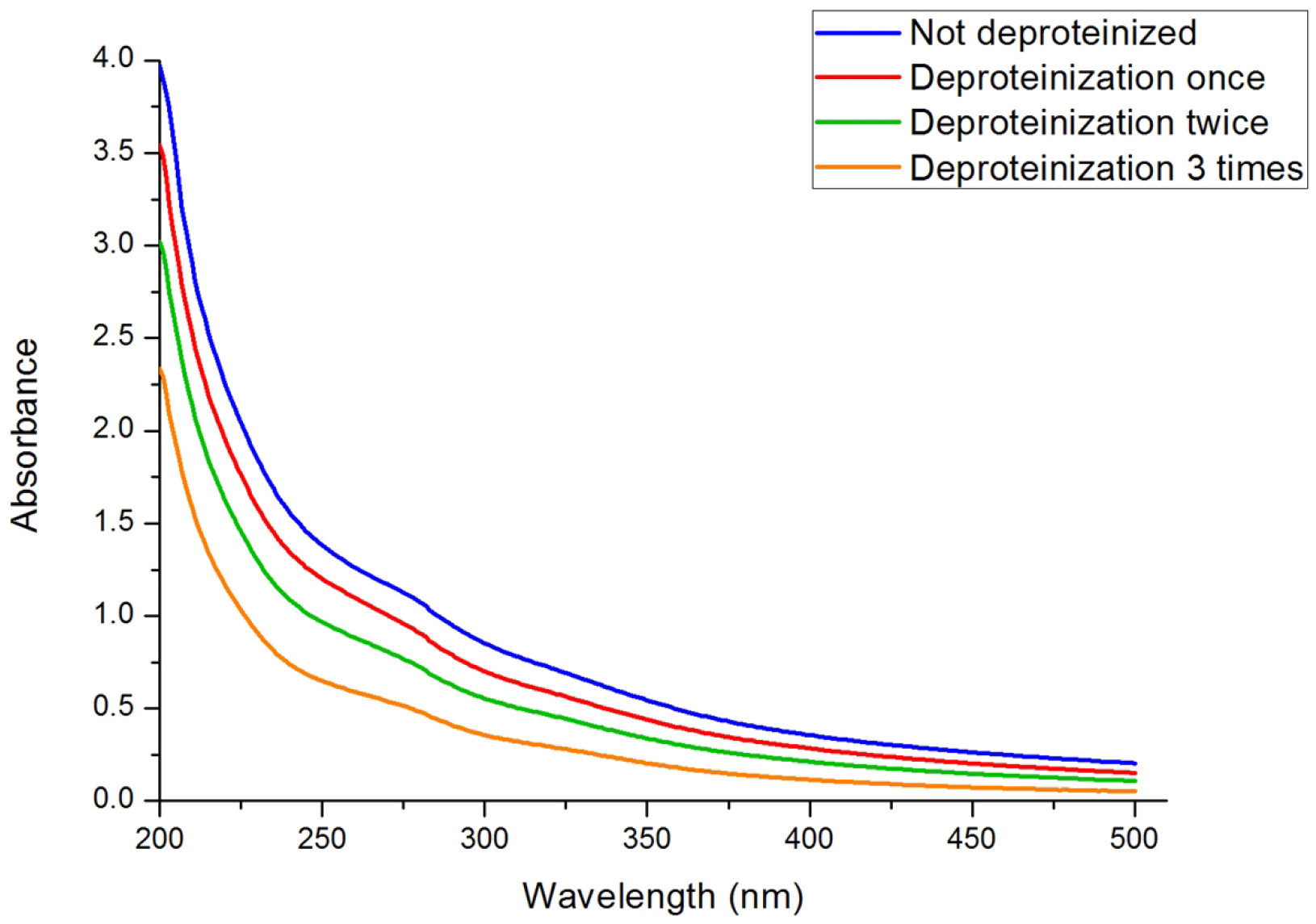
UV spectrum of rice bran polysaccharide

#### 3.2.3 ^13^C NMR

At present, we want to find out the connection position between monosaccharide, and the commonly used method is nuclear magnetic resonance detection. A sample of 50 mg rice bran polysaccharide was dissolved in 0.5 mL D_2_O.

After loading the sample, we tested it at 20°C at room temperature. After 3 hours, we got the results as shown in **Fig.9.** We can infer that the connecting end is α-D-glycosidic bond structure; The attribution of the peaks is as follows:62.51ppm(C-6);71.21ppm(C-5);73.37ppm(C-2);76.75ppm(C-4);78.75ppm(C-3);99.92ppm(C-1). The above indicates that rice bran polysaccharide (RBPs) has two glycosidic bond forms of α-1, 4 and α-1, 6, of which α-1, 4 has a large structural content.

**Fig.9.**
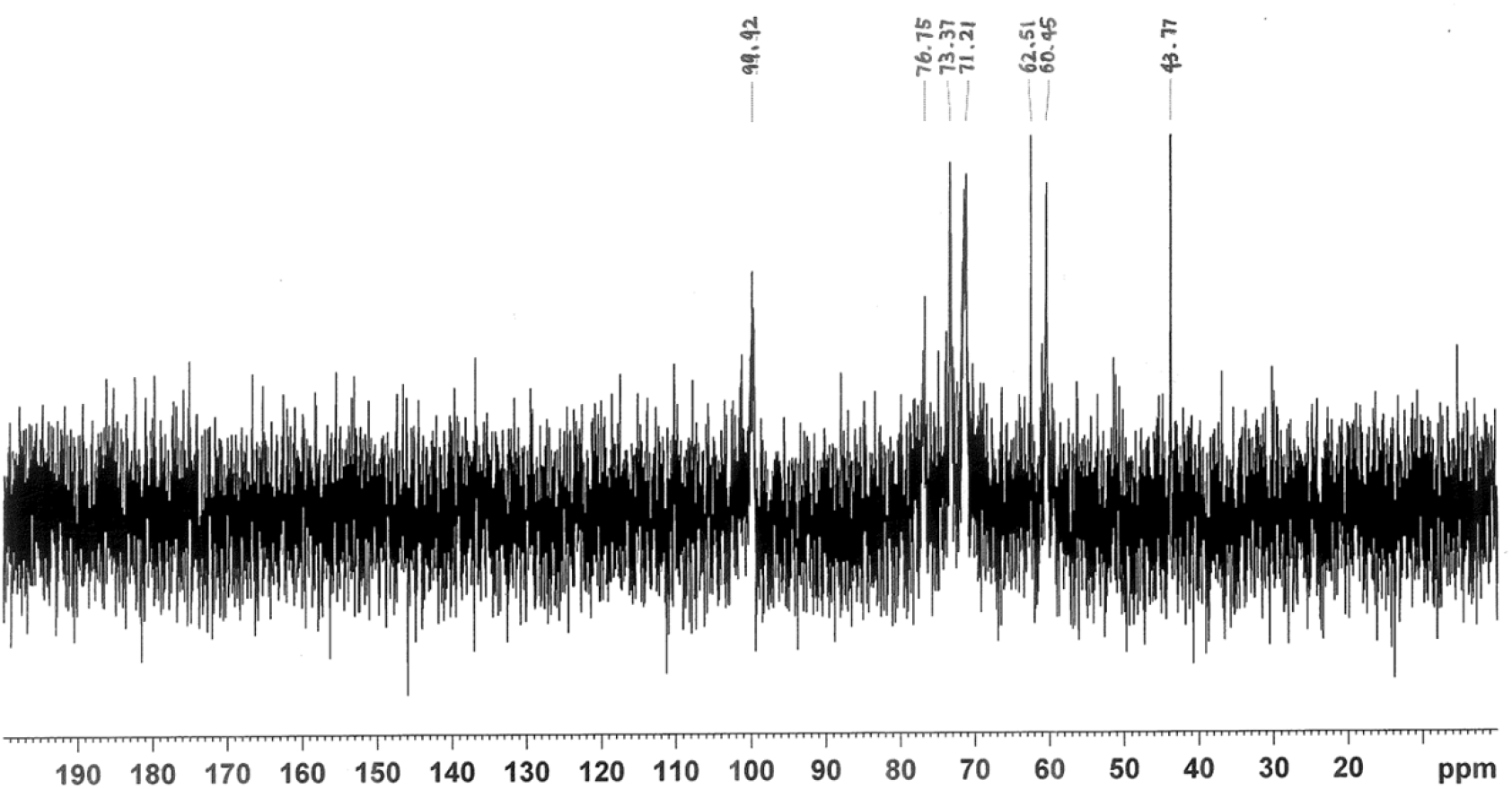
^13^C NMR of rice bran polysaccharide

**Fig.10.**
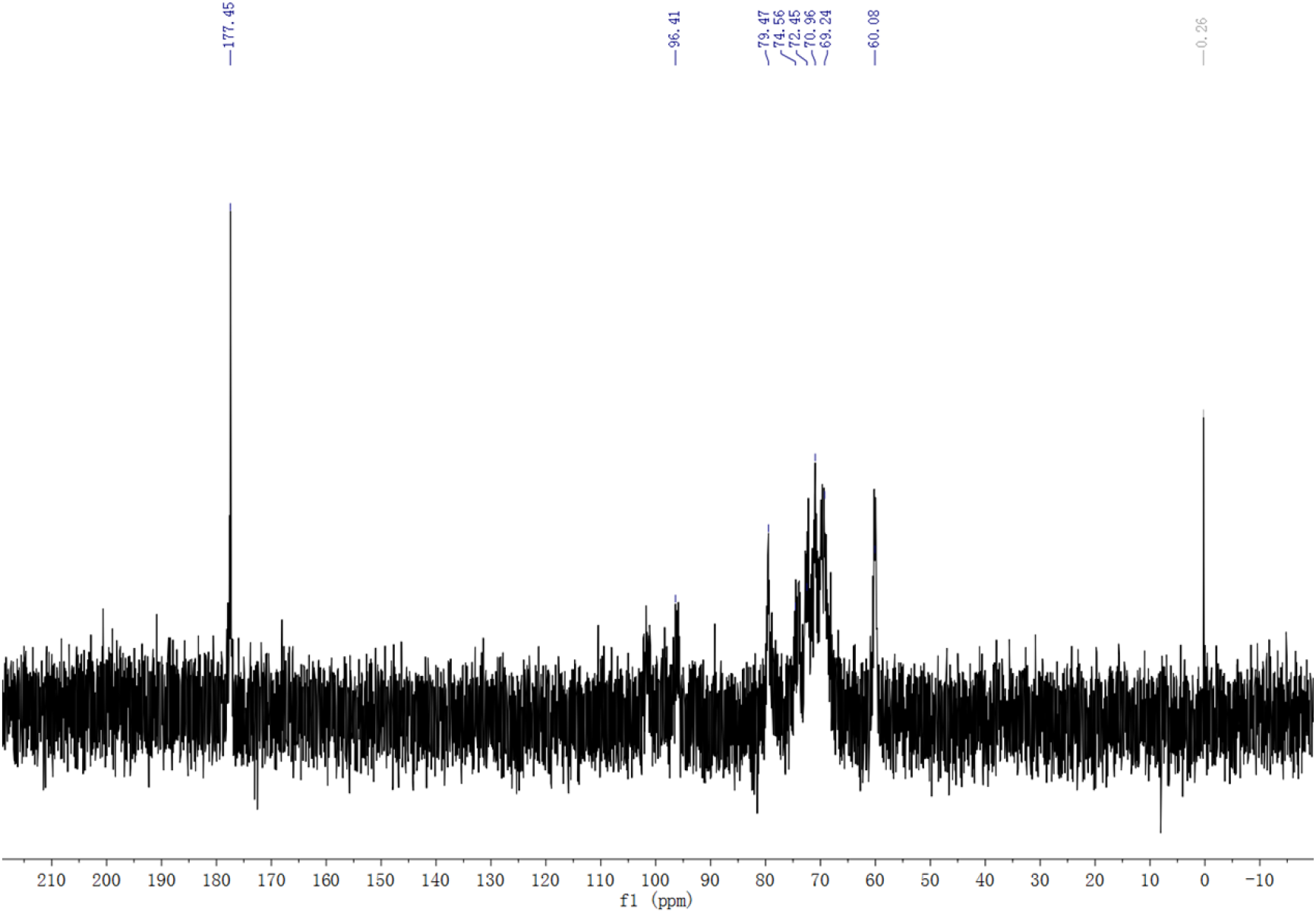
13C NMR of carboxymethylated rice bran polysaccharide

**Fig.11.**
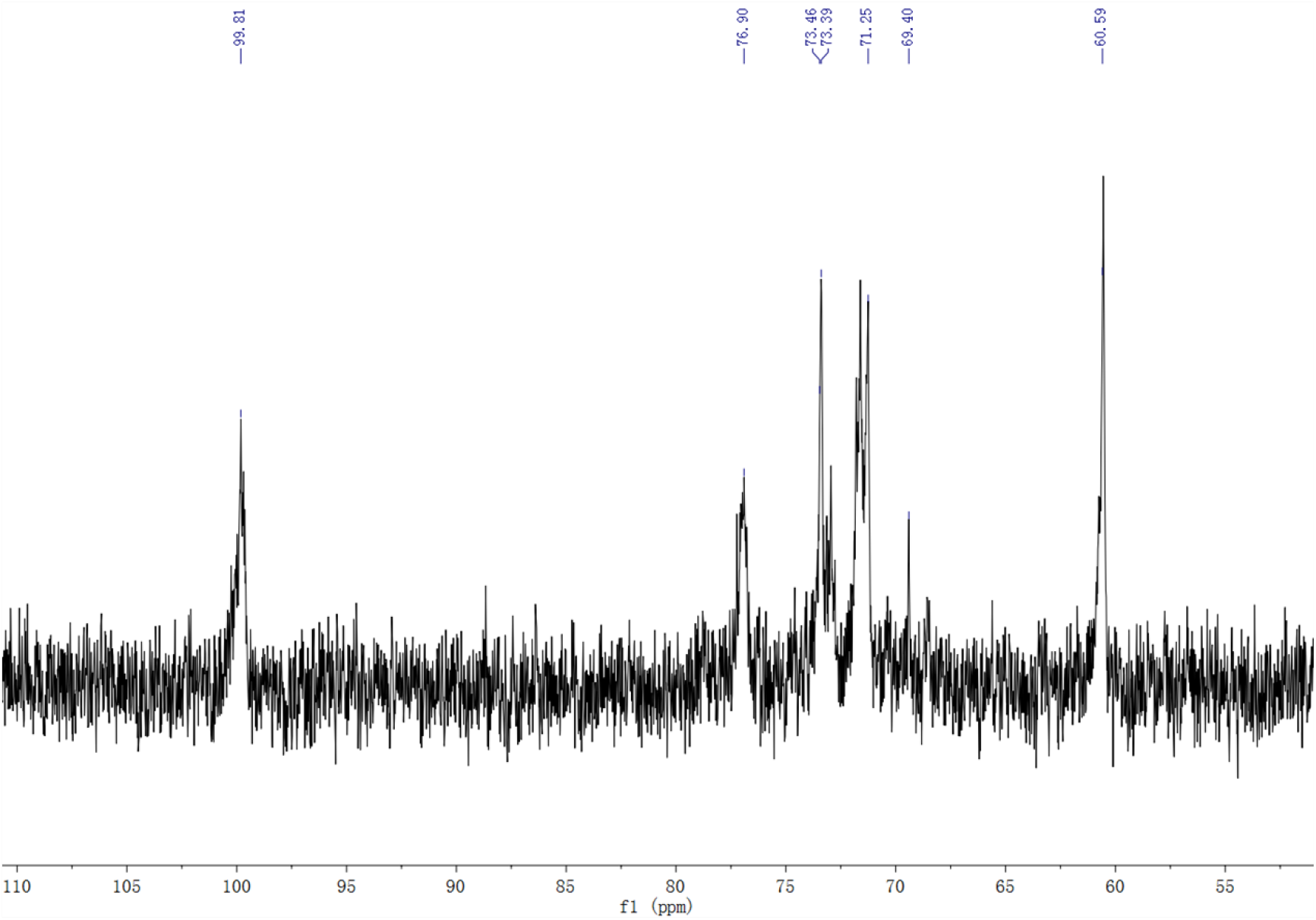
^13^C NMR of phosphorylated rice bran polysaccharide

At present, to find out the connection position between monosaccharides, the common method is nuclear magnetic resonance detection. A sample of 50 mg carboxymethylated rice bran polysaccharide was dissolved in 0.5 mL D_2_O. We can infer that the connecting end is α-D-glycosidic bond structure; The attribution of the peaks is as follows :60.08ppm(C-6);69.24ppm(C-5);2.45ppm(C-2);74.55ppm(C-4); 79.47ppm(C-3);96.41ppm(C-1);The new chemical shifts at 177.45ppm could be attributed to the (C-O) of carboxymethyl group. The above indicates that rice bran polysaccharide (RBPs) has two glycosidic bond forms ofα-1, 4 andα-1, 6, of whichα-1, 4 has a large structural content.

A sample of 50 mg phosphorylated rice bran polysaccharide was dissolved in 0.5 mL D_2_O.After loading the sample, we tested it at 20°C at room temperature. After 3 hours, we got the results as shown in **Fig.11.**

99.81ppm(C-1);73.39ppm73.46ppm(C-2);68.56ppm(C-4);76.90ppm(C-5);60.59 ppm(C-6)71.25ppm68.4ppm(C-7);

A sample of 50 mg acetylated rice bran polysaccharide was dissolved in 0.5 mL D_2_O.After loading the sample, we tested it at 20°C at room temperature. After 3 hours, we got the results as shown in **Fig.12.**

**Fig.12.**
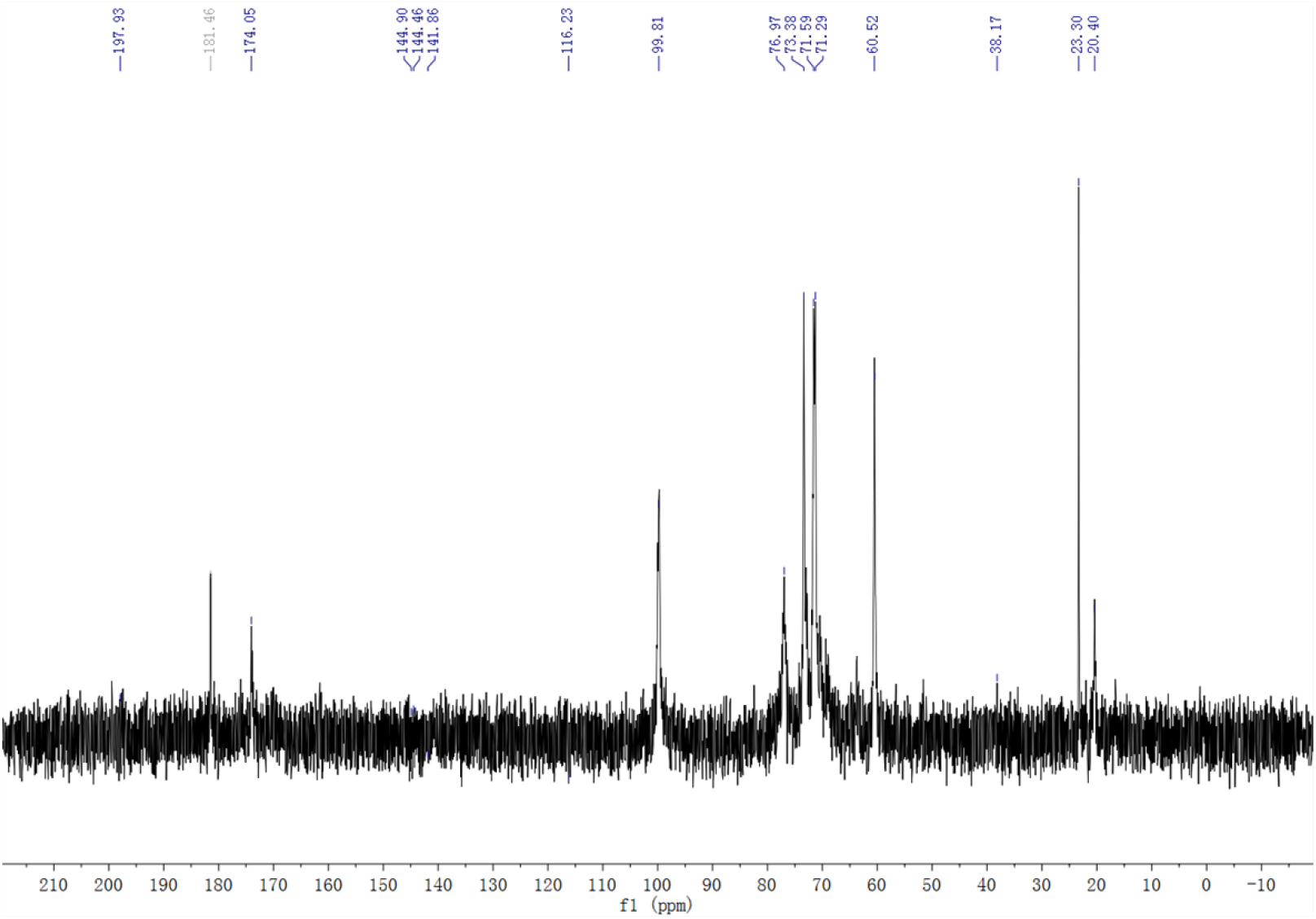
^13^C NMR of acetylated rice bran polysaccharide

99.81ppm(C-1);60.52ppm(C-6);71.29,71.59ppm(C-3);73.38ppm(C-2);76.97ppm (C-5);174.05ppm(C-8);181.46ppm(C-9)

## 4 Various antioxidant activities

### 4.1 Determination of hydroxyl radical scavenging ability

Reaction mechanism: Fe^2+^+H_2_O_2_==·OH+OH^−^+Fe^3+^, as shown in **Fig.13.**

**Fig.13.**
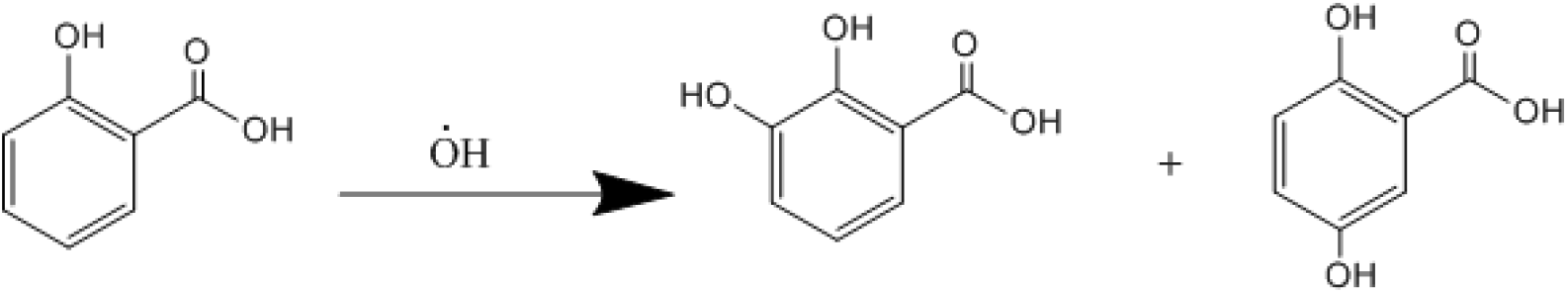
Reaction principle of hydroxyl radical with salicylic acid

Salicylic acid can react with it to generate 2,3-dihydroxybenzoic acid and 2,5-dihydroxybenzoic acid. These two isomers have UV absorption at 510nm. The stronger the absorption, it indicates that the salicylic acid is bound. The more hydroxyl radicals you have. Vc or polysaccharide can also scavenge hydroxyl radicals. If the same amount of salicylic acid is added, the same hydroxyl radicals are generated. The ability of polysaccharides to scavenge hydroxyl radicals is determined by measuring the difference in absorption at 510 nm after adding polysaccharides. As shown in **Fig.14.**

**Fig.14.**
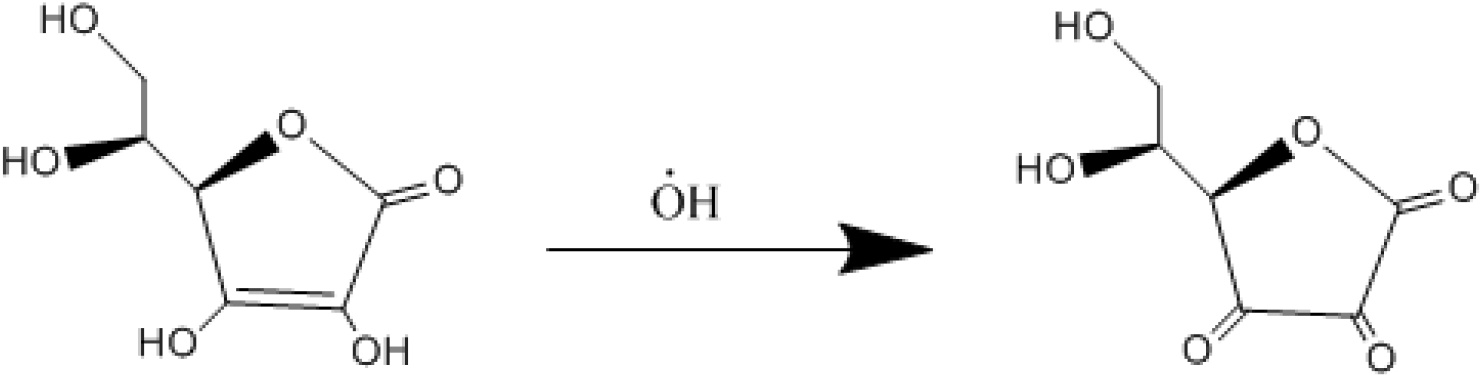
Mechanism of V_c_ scavenging hydroxyl radicals

Specific experimental steps: The sample solution (1mL) of rice bran polysaccharide and its derivatives with concentrations of 0.1, 0.2, 0.4, 0.8, 1.6, 3.2 mg/mL was added to the test tube(distilled water was used as a control).1mL 9×10^−3^mol/L FeSO_4_, 1mL 9×10^−3^mol/L salicylic acid-ethanol solution (70% aqueous solution) were added to the test tube.After being thoroughly mixed, 9×10^−3^mol/L H_2_O_2_ solution (1mL) was added to the test tube.They were all placed in a constant temperature oil bath at 37°C and reacted for 30 minutes.After the solution was cooled to room temperature, the UV absorbance of the solution was measured at 510 nm and recorded as As. Vc was used as a positive control group, and the test was repeated three times.

The hydroxyl radical scavenging rate E is calculated according to the following formula:

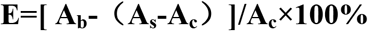

**A**_**b**_: **Substitute the absorbance of polysaccharide sample with distilled water**

**A**_**s**_: **Absorbance after adding the sample solution**

**A**_**c**_: **Absorbance of rice bran polysaccharide or its derivative solution itself**

Vc was used as a positive control group of rice bran polysaccharides and their derivatives to study the ability of rice bran polysaccharides and their derivatives to scavenge hydroxyl radical ions.As shown in **Fig.15**, rice bran polysaccharides and several derivatives at a concentration of 0.1-3.2mg/mL showed the ability to scavenge •OH with increasing concentration. Among them, the ability of low-concentration Ac-RBPs (0.1-0.2mg/mL) to remove •OH is higher than that of the same concentration of RBPs, but with the concentration increases, the ability of Ac-RBPs to remove •OH is significantly lower than RBPs. However, both P-RBPs and CM-RBPs significantly improved the ability to remove •OH. Among them, the P-RBPs at a concentration of 3.2 mg/mL showed the best ability to remove •OH, with a removal rate of 68.9%. Compared with the same concentration of rice bran polysaccharide, the scavenging rate of hydroxyl radicals (27.8%) increased by 41.1%.But they are still lower than Vc’s ability to remove •OH.

**Fig.15.**
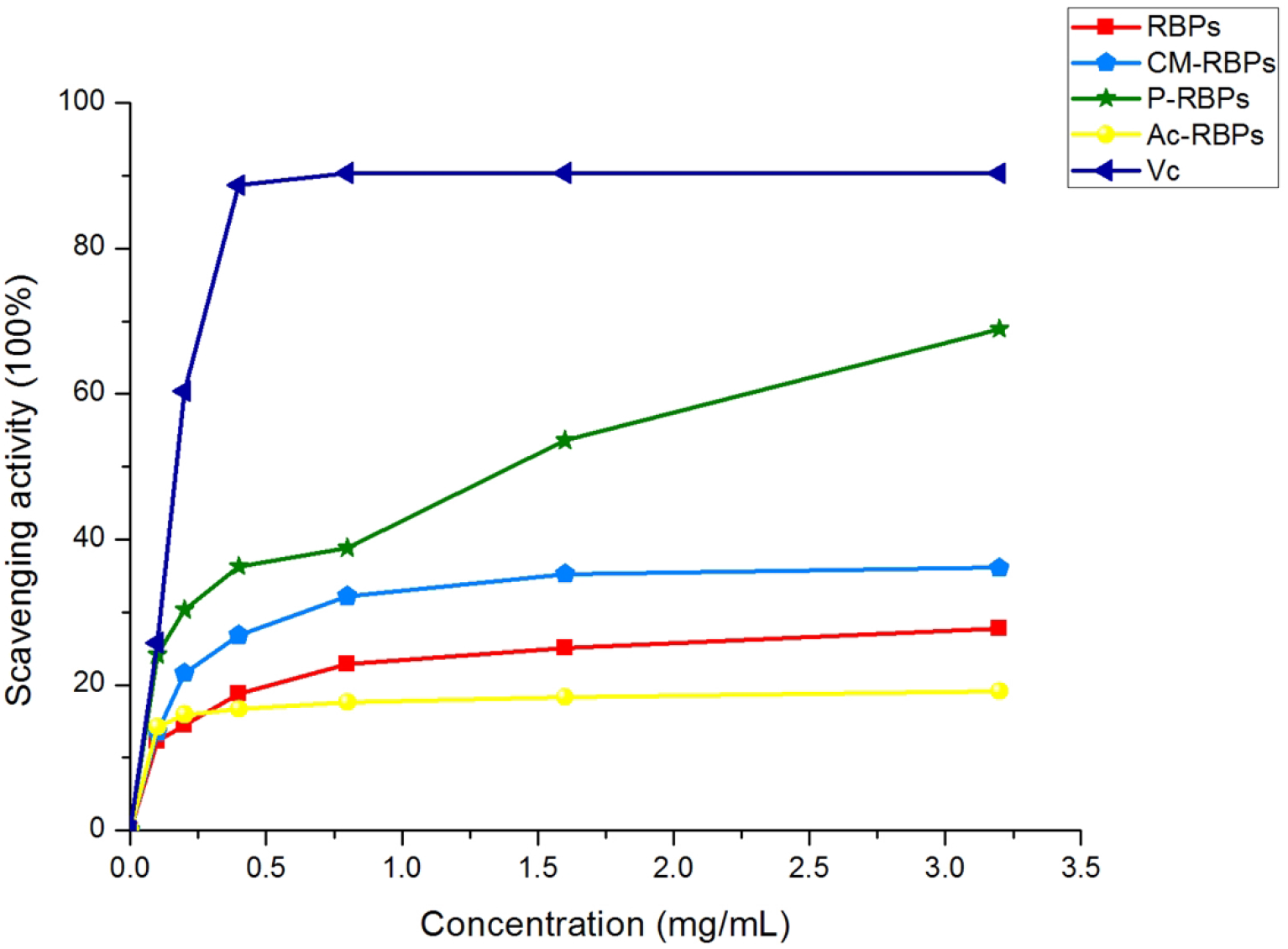
Results of hydroxyl radical ion scavenging ability

### 4.2 Superoxide anion scavenging ability

In an environment with a pH value greater than 7, pyrogallic acid can reflect its strong reducing ability and undergo an auto-oxidation reaction, simultaneously generating superoxide anion radicals (O_2_^−^•) and colored intermediates. The reaction mechanism is shown in **Fig.16** The intermediate product has a characteristic ultraviolet absorption peak at λ = 320nm. In the early stage of the reaction, the total amount of intermediate products is positively correlated with time. When added with scavengers such as Vc to remove superoxide anions, it can react with O_2_^−^•, which can effectively prevent the continued formation and accumulation of intermediate products, resulting in a weakened UV absorption of the solution at λ = 320nm. Therefore, the ability of rice bran polysaccharide to remove O_2_^−^• can be judged by measuring the value there.

**Fig.16.**
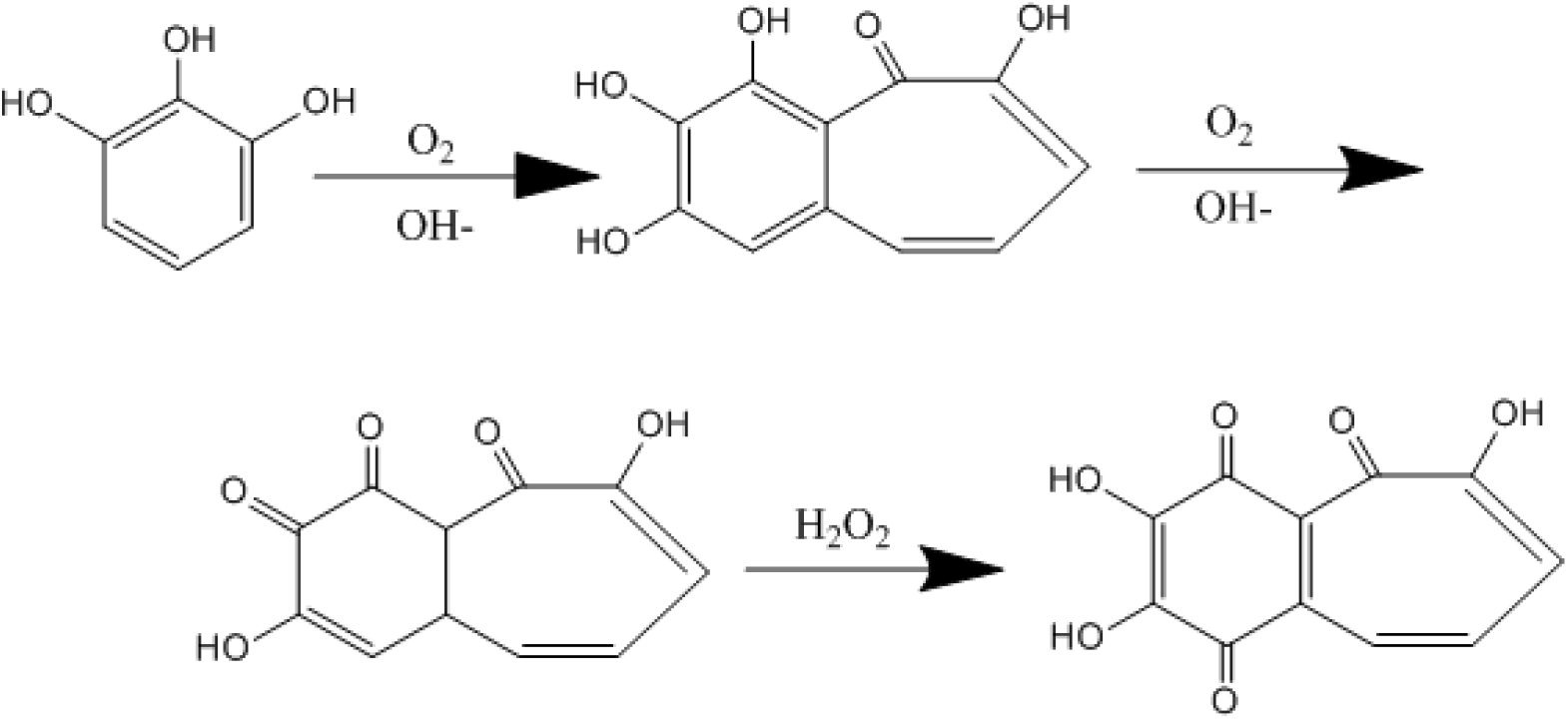
Pyrogallol auto-oxidation process

Specific experimental steps: The concentration of 0.1, 0.2, 0.4, 0.8, 1.6, 3.2 mg/mL rice bran polysaccharide and its derivative solutions (0.2 mL) were added to different test tubes (the control group was replaced with distilled water).3 mL of 0.05mol/L, pH 8.2 Tris•HCl buffer solution was added to the test tube. They were all placed in a constant temperature oil bath with a temperature of 25°C, and holding 10 minutes. The preheated 30mmol/L pyrogallic acid solution (12uL) was added to the test tube, and reacted accurately for 4 minutes. The reaction has been terminated with 0.5mL concentrated hydrochloric acid. The absorbance at 320nm was measured.And use an equal volume of pH 8.2 Tris • HCl buffer solution as a blank control group, and Vc as a positive control group, the test was repeated three times.

The hydroxyl radical scavenging rate E is calculated according to the following formula:

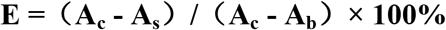

**A**_**b**_: **Tris • HCl buffer absorbance**

**A**_**s**_: **Absorbance after adding the sample solution**

**A**_**c**_: **Tris • HCl buffer solution + absorbance of pyrogallic acid**

Vc was used as a positive control group of rice bran polysaccharide and its derivative solution to study the ability of rice bran polysaccharide and its derivatives to remove superoxide anions. From the **Fig.17**, it can be found that Ac-RBPs shows relatively good ability to remove superoxide anions.At a concentration of 3.2 mg/mL, the superoxide anion scavenging rate of Ac-RBPs reached 49.9%.That scavenging rate of rice bran polysaccharide at low concentration (0.1-0.8mg/mL) is lower than that of the other two derivatives.However, the scavenging rate of rice bran polysaccharides at high concentrations (1.6-3.2mg/mL) to superoxide anions is even slightly higher than that of the other two derivatives.However, the scavenging ability of superoxide anion for all sample groups is still less than Vc.

**Fig.17.**
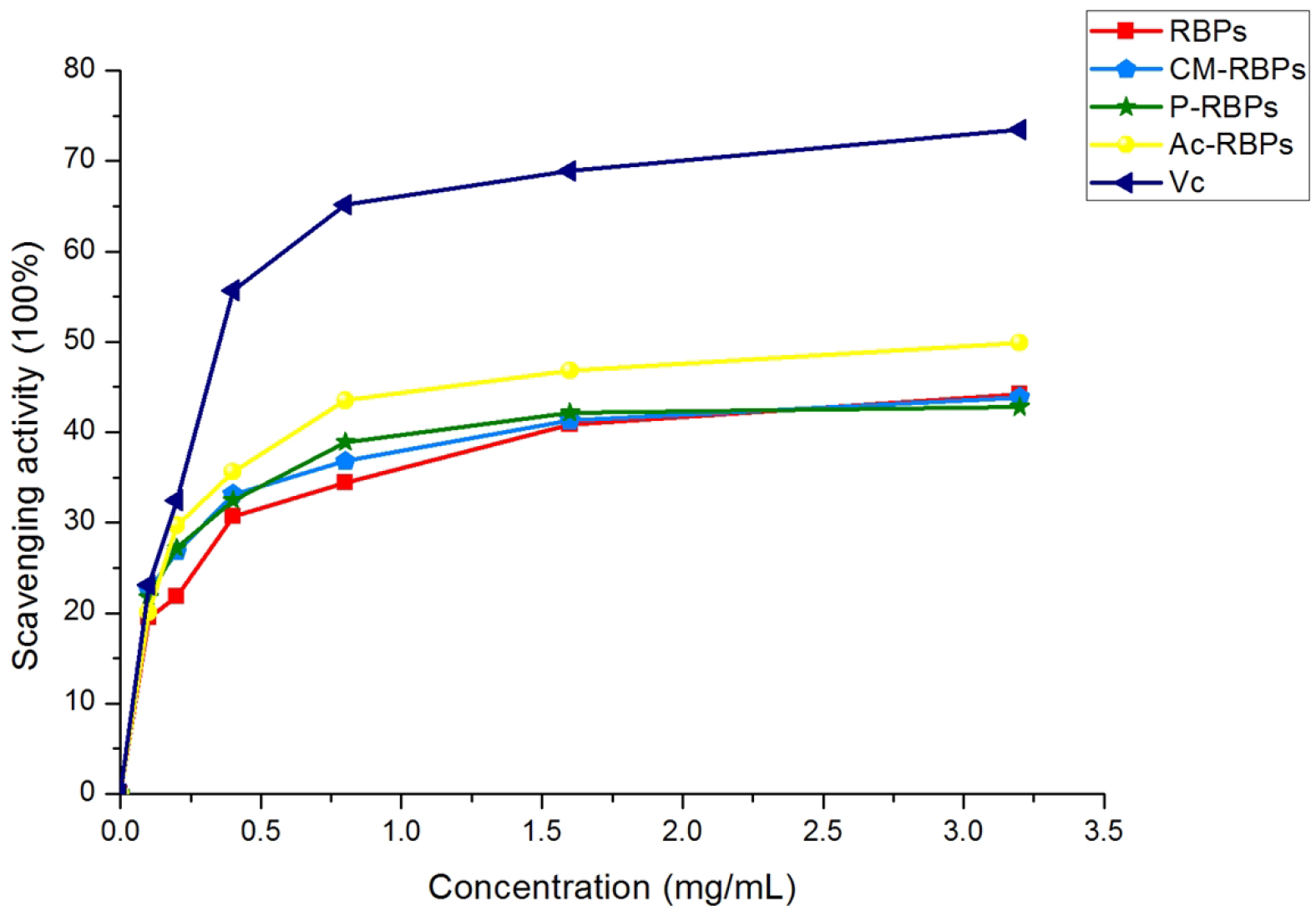
Results of superoxide anion scavenging ability

### 4.3 Reduction ability

Analysis of reducing power measures the electron-donating ability of antioxidants using the potassium ferricyanide reduction method [30].The reducing activities were usually related to the development of reductones, which terminate free radical chain reactions by donating a hydrogen atom[31]. The **Fig.18** reflects the principle of detecting the reducing ability of polysaccharides.

**Fig.18.**
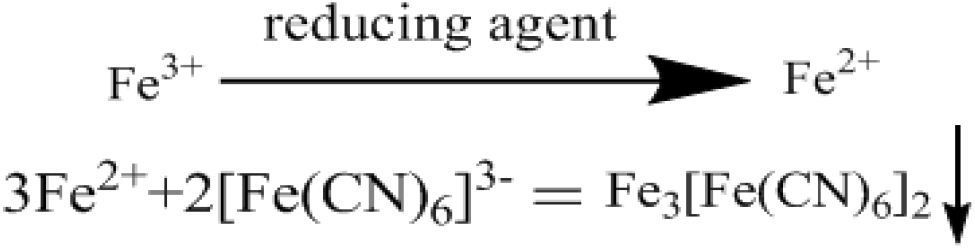
Reaction mechanism

The samples with different concentrations (0.1,0.2,0.4,0.8,1.6,3.2 mg/mL,1.0 mL) were mixed with 2.5 mL of PBS (pH 6.6) and 2.5 mL of potassium ferricyanide (1%).The mixture was incubated for 30 min at 50 ◦C. The reaction was termi-nated by TCA solution (10%).Then, the solution (2 mL) was mixed with distilled water (2 mL) and ferric chloride (0.1%, 0.4 mL).Measure its UV absorbance at 700 nm, the higher the absorbance, the higher the reduction ability. Vc was used as a control group.

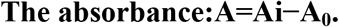

**Ai:The absorbance of sample.**

**A**_**0**_:**The absorbance of control.**

It can be seen from **Fig. 19** that the reducing ability of several samples increases with the increase of concentration.The reducing ability of rice bran polysaccharides is significantly weakened after carboxymethylation and acetylation. Compared with the same concentration of rice bran polysaccharides, there is almost no difference in reducing ability of phosphorylated rice bran polysaccharides. Several derivatizations are not helpful in improving the reducing ability of rice bran polysaccharides.The reducing ability of rice bran polysaccharide and its derivatives is much weaker than that of Vc.

**Fig.19.**
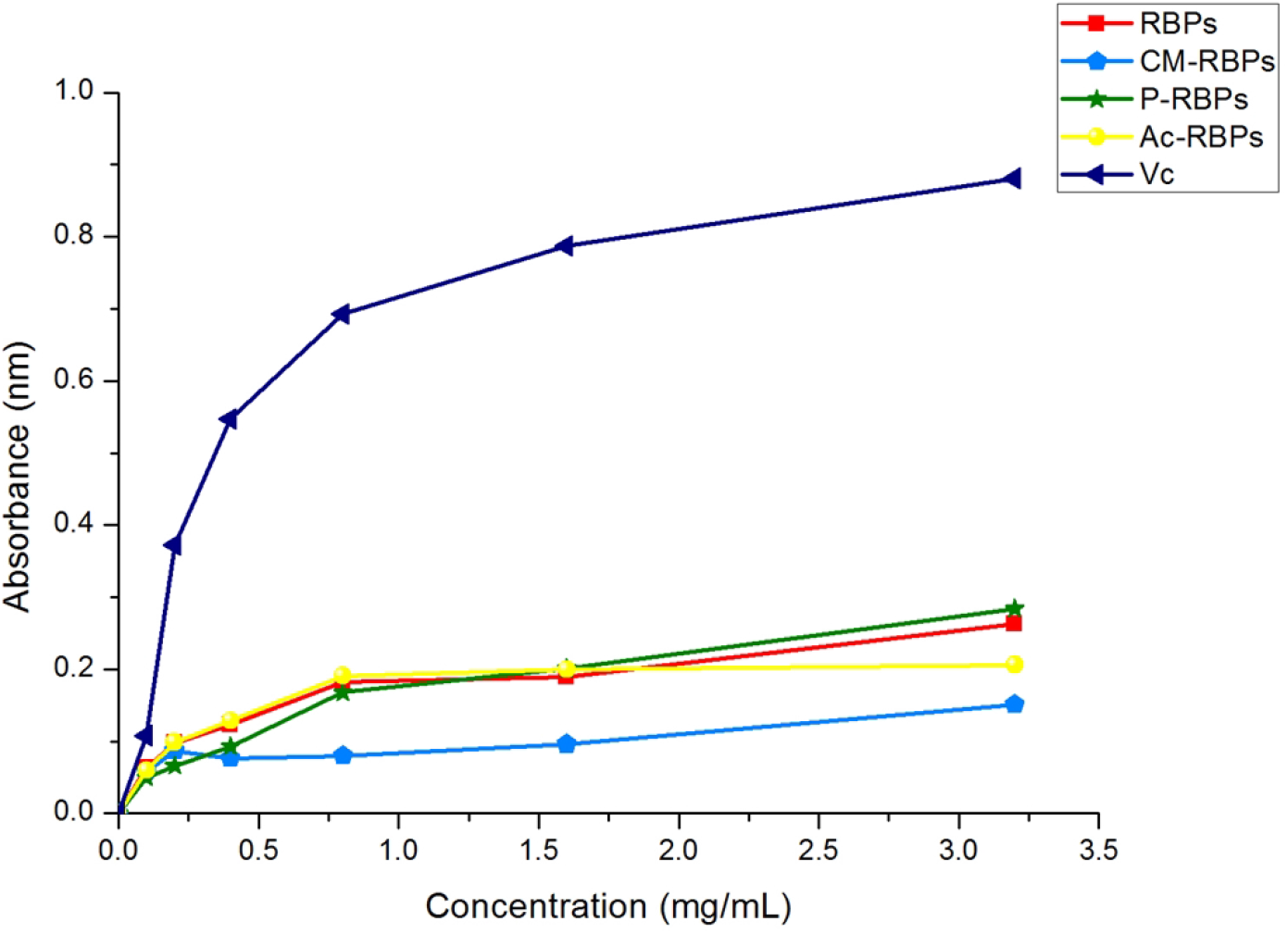
Results of reduction ability

### 4.4 Determination of anti-lipid peroxidation ability

0.1g of soy lecithin has been weighed. Soy lecithin has been mixed with pH = 7.4 PBS (100 mL). The rice bran polysaccharide solution or its derivative solution (4 mL) has been added to each test tube at a concentration of 0.1, 0.2, 0.8, 0.4, 1.6, 3.2 mg/mL. And soy lecithin solution (3.6mL), 10mmol/L FeSO_4_ (0.4 mL) solution, 10mmol/L Vc solution (0.4 mL), have been added to the test tube.Incubate at 37°C for 60 min and cool to 25°C. Add 1.0 mL of trichloroacetic acid (TCA, 20%, w/v) and 1.0 mL of thiobarbituric acid (TBA, 0.8%, w/v) successively, and heat the mixture at 100°C for 15 minutes. Centrifuge at 3000 rpm Collect the supernatant of the sample after 15 minutes. The discoloration was measured at 532 nm.

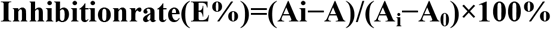

**A**_**0**_:**The absorbance value of Tris-HCl buffer.**

**A**_**i**_:**The absorbance value of Tris-HCl buffer solution and pyrogallol solution.**

**A: The absorbance of sample solution.**

According to **Fig. 20**, the anti-lipid peroxidation ability of carboxymethylated rice bran polysaccharide, phosphorylated rice bran polysaccharide, and acetylated rice bran polysaccharide all exceeded the ability of underivatized rice bran polysaccharide. Of particular concern is the anti-lipid peroxidation ability of high concentration (3.2mg/mL) phosphorylated rice bran polysaccharide even surpasses that of the same concentration of Vc.

**Fig.20.**
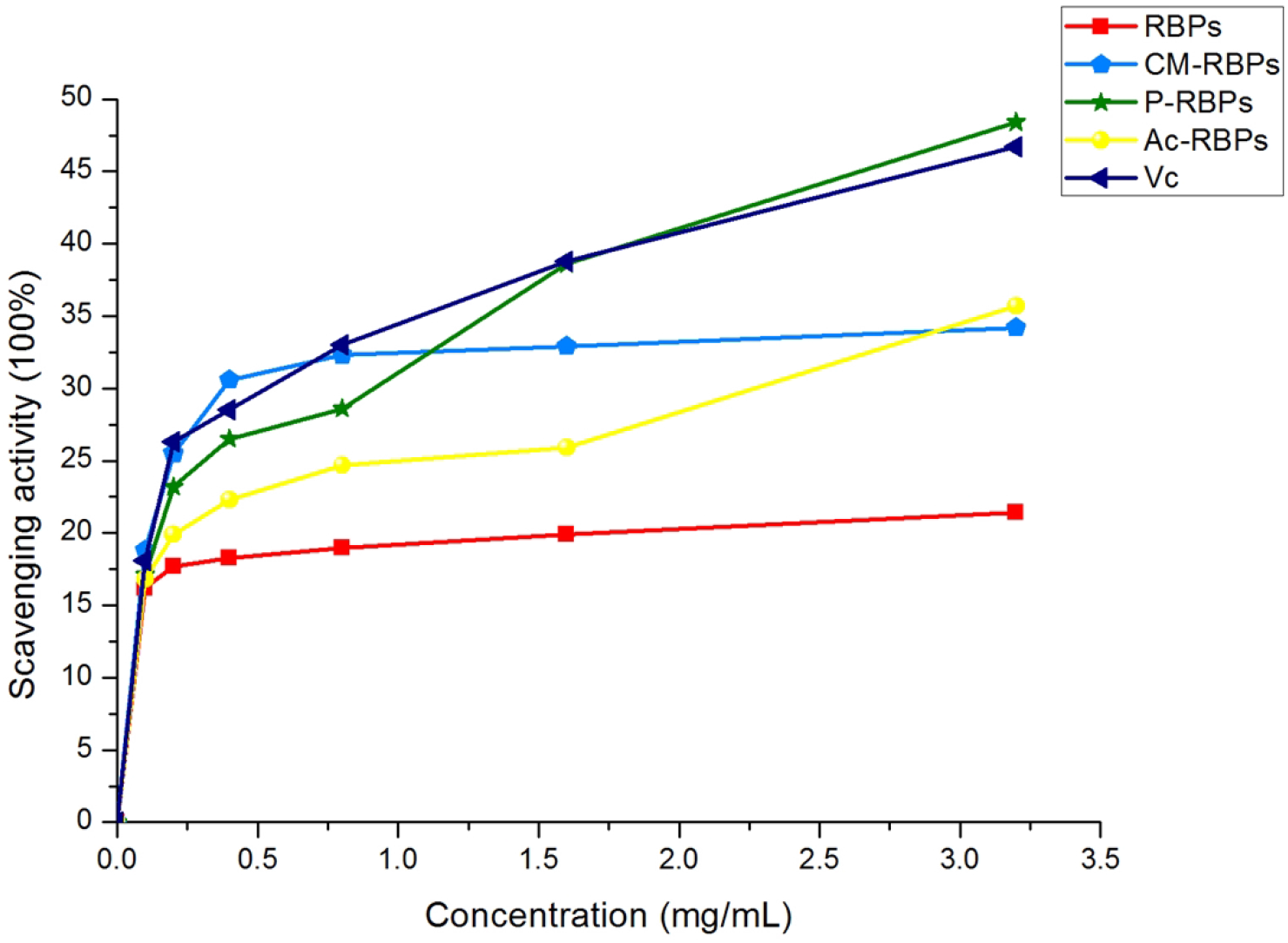
Results of anti-lipid peroxidation

## 5 Results and discussion

In addition to further proof of its polysaccharide composition, infrared characteristic absorption spectra at 761, 848, and 933 cm−1 were also found in the fingerprint area. These results can prove that the sample was composed of α-D-glucopyranose. In addition to further proof of the infrared results, a carbon spectrum nuclear magnetic resonance method was used to find the glycosidic bonds between the glycan rings: the connection mode of α-1,4 and α-1,6 glycosidic bonds. And found that the anti-lipid peroxidation ability of phosphorylated rice bran polysaccharide is better than Vc. In the next work, we are also going to carry out carboxymethyl phosphorylation, sulfation and carboxymethyl sulfation of rice bran polysaccharides and further study the antioxidant activity of rice bran polysaccharide derivatives. In order to prepare derivatives with better antioxidant activity.

